# Global variation in the ratio of sapwood to leaf area explained by optimality principles

**DOI:** 10.1101/2024.02.24.581904

**Authors:** Huiying Xu, Han Wang, I. Colin Prentice, Sandy P. Harrison, Lucy Rowland, Maurizio Mencuccini, Pablo Sanchez-Martinez, Pengcheng He, Ian J. Wright, Stephen Sitch, Qing Ye

**Affiliations:** Department of Earth System Science, Ministry of Education Key Laboratory for Earth System Modeling, Institute for Global Change Studies, Tsinghua University, Beijing 100084, China; Department of Geography, University of Exeter, Exeter, EX4 4RJ, UK; Georgina Mace Centre for the Living Planet, Department of Life Sciences, Imperial College London, Silwood Park Campus, Buckhurst Road, Ascot, SL5 7PY, UK; School of Archaeology, Geography and Environmental Sciences (SAGES), University of Reading, Reading, RG6 6AH, UK; CREAF, Campus UAB, Cerdanyola del Vallés 08193, Spain; ICREA, Barcelona 08010, Spain; Universitat Autònoma de Barcelona, Cerdanyola del Vallés, Barcelona 08193, Spain; South China Botanical Garden, Chinese Academy of Sciences, Xingke Road 723, Guangzhou 510650, China; Hawkesbury Institute for the Environment, Western Sydney University, Penrith, New South Wales, Australia; School of Natural Sciences, Macquarie University, North Ryde, NSW 2109, Australia

## Abstract

The sapwood area supporting a given leaf area (*v*_H_) reflects a coordinated coupling between carbon uptake, water transport and loss at a whole plant level. Worldwide variation in *v*_H_ reflects diverse plants strategies adapt to prevailing environments, and impact the evolution of global carbon and water cycles. Why such a variation has not been convincingly explained yet, thus hinder its representation in Earth System Models. Here we hypothesize that optimal *v*_H_ tends to mediate between plant hydraulics and leaf photosynthesis so that leaf water loss matches water supply. By compiling and testing against two extensive datasets, we show that our hypothesis explains nearly 60% of *v*_H_ variation responding to light, vapour pressure deficit, temperature, and sapwood conductance in a quantitively predictable manner. Sapwood conductance or warming-enhanced hydraulic efficiency reduces the demand on sapwood area for a given total leaf area and, whereas brightening and air dryness enhance photosynthetic capacities consequently increasing the demand. This knowledge can enrich Earth System Models where carbon allocation and water hydraulics play key roles in predicting future climate-carbon feedback.

## Introduction

Tightly coupled terrestrial carbon and water cycling that influence global energy balance and climate system are mediated by complex plant physiological processes and traits. Multiple plant processes and traits interact with one another, but regulate plant performance in response to fluctuating environment at different time scale. Leaf photosynthesis and transpiration are closely linked, as stomata guard the gate of water loss and CO_2_ absorption at the same time. Leaf photosynthesis is affected by intrinsic plant properties, such as maximum photosynthetic capacities. Meanwhile, the plant hydraulic properties influence transpiration. These plant processes and traits are coordinated, resulting in the balance of carbon allocation to different functional organs (i.e. ratio of sapwood to leaf area). Despite the coupled trait network, photosynthesis-related traits can vary at relatively short timescale under changing environment compared to plant hydraulic traits. The hydraulic properties that often related to wood/leaf anatomy shifts at yearly timescale, thus species turnover might involve in their response to climate change at a community level. Over recent decades there has been continuous atmospheric warming and rising vapour pressure deficit (*D*) worldwide, leading to increased drought and heatwave frequency and severity, severely impacting plant functioning (Trenberth et al. 2013; Dai et al. 2018; Fu et al. 2022). It is pertinent to investigate how plant strategies adapt/acclimate to climate so that we can improve our understanding of carbon and water cycling under global climate change.

Diverse plant eco-physiological traits and allocation strategies in turn are shaped by the surrounding environments, reflecting various adaptation strategies evolved through natural selection. Eco-evolutionary optimality principles (EEO) offers an alternative perspective to examine the effect of environment on plant strategies based on the idea that plants adapt to their surrounding environment through evolutionary processes (Franklin et al., 2020; Harrison et al., 2021). It has been successfully applied to explain optimal trait behaviour in response to climate, such as maximum capacity of carboxylation (*V*_cmax_), ratio of leaf-internal to ambient CO_2_ partial pressure (χ) and leaf mass per area. However, previous applications of EEO mainly focus on leaf-level traits. Seldom studies consider physiological processes across different organs due to complicated and ambiguous trait coordination networks. The xylem hydraulic conductivity (*K*_S_) has long been observed to be related to wood density and scale with the ratio of sapwood to leaf area (*v*_H_) to maintain sufficient water supply. Furthermore, there is evolutionary correlation between *K*_S_ and *v*_H_, suggesting their co-evolution under selective environment (Sanchez-Martinez et al., 2020). Leaf water potential at turgor loss point (Ψ_tlp_) is close to minimum water potential and stomatal closure point, which indicates its control on photosynthesis and stomatal behaviour (Zhu et al., 2017). The difference between leaf and soil water potential drives constant water flow. Hence, the hydraulic, photosynthetic traits and allocation strategy are tightly coupled at individual level but less explored. Recent study shows that *v*_H_ sits at the centre of trait coordination networks and mediates multiple processes in the coupling of plant water-carbon cycle. Whether or not its variation and its trade-off with other hydraulic traits could be explicitly explained by a unified framework of optimal regulation on the carbon-water coupling hasn’t been tested globally.

Due to the lack of understanding on the variation in *v*_H_, process-based models as a tool of simulating the future earth system show large uncertainties in their predictions on future climate-carbon feedback, especially under drought and heatwave events. Current representation of carbon allocation is either derived from empirical relationships involving response to environment without considering its coordination with other physiological processes or plenty of tuneable plant functional types parameters. A realistic, efficient and unified model of *v*_H_ that can be directly implemented in process-based models is urgently needed to improve land-surface models (LSMs).

Here based on EEO concepts, we propose a universal parsimonious theory to explain the variation in *v*_H_ as a trade-off with hydraulic traits regulated by environment in order to coordinate photosynthesis, transpiration and hydraulics at whole-plant level. In other words, the water demand by transpiration for supporting canopy photosynthesis equals the water supply from trunk water transport. The environmental variables (irradiance, temperature, vapour pressure deficit and CO_2_) drive those processes and consequently shape the variation in *v*_H_ and its trade-off with hydraulic traits. We test our theory using two global hydraulic traits datasets collected at species and individual-at-site levels. We show that our theory predicts *v*_H_ variation at global scale. This model can be incorporated into LSMs as flexible plant allocation and hydraulic schemes to improve the prediction of land carbon and water cycling under future climate.

## Theory

Based on EEO, we hypothesize that maximum plant water transport through the xylem matches maximum water demand required to maintain photosynthesis, considering resources are optimally allocated (Whitehead et al. 1984). In other words, the water uptake from soil driven by the maximum water potential difference between soil and leaf (ΔΨ_max_) is transported through root, sapwood to leaf evaporating surface for transpiration through stomata so that in turn CO_2_ from atmosphere is absorbed to sustain optimal photosynthetic capacity.

The canopy water demand is determined by canopy total leaf area and transpiration rate per leaf area. The latter factor is regulated by stomata, and predictable from EEO-based least-cost and coordination hypotheses via the coupling between photosynthesis and transpiration using Fick’s law. The carbon assimilation is co-limited by carboxylation and electron transport processes. The coordination hypothesis states that light- and Rubisco-limited photosynthetic rates should be equal to optimally utilize light without extra carbon costs (Wang et al., 2017). The leaf internal CO_2_ level for carboxylation is controlled by stomatal conductance, and optimal ratio of leaf-internal to ambient CO_2_ (χ) is achieved by minimizing unit carbon cost of photosynthesis and transpiration over weeks to months (Prentice et al., 2014). Both traits determine the maximum water demand for leaf gas exchange and optimal photosynthesis. Further combing with the classic biochemical model of photosynthesis and Fick’s law, the leaf level transpiration demand can be predicted as a function of irradiance, temperature, vapour pressure deficit (*D*) and CO_2_. The trunk water supply is dominated by hydraulic properties, including compensation effect of maximum hydraulic efficiency and driving force of ΔΨ_max_. The maximum hydraulic efficiency is influenced by structure of hydraulic systems (i.e. pit membrane, conduit diameter), and temperature via water viscosity and cell membrane permeability. All else being equal, plants with higher hydraulic efficiency (due to enhanced *K*_S_ or warming-reduced water viscosity and cell membrane permeability) can transport more water through a given stem, which could increase total leaf area. We assumed that (1) this match between water supply and demand often occurs around noon where stomatal conductance and transpiration reach their peak; (2) the optimal *v*_H_ is achieved at multiple-year timescale under environmental selection. Although photosynthetic traits optimize at weekly to monthly timescale, the hydraulic properties, especially maximum hydraulic efficiency, have limited plasticity. *v*_H_ is the intermediate trait to coordinate both sets of traits in respond to the surrounding environment and optimized to meet the balance as a result of coordinated physiological processes (Xu et al. 2021). At site level, the optimal *v*_H_ is achieved through plasticity and species turnover over years.

These processes eventually lead to an optimal *v*_H_ balancing between the hydraulic efficiency in trunks and the water demand from canopy as predicted by climates of irradiance, temperature and *D*:

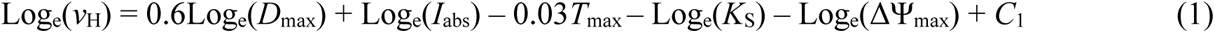

Here *C*_1_ is the parameter containing information about photosynthetic traits values under standard climate condition and plant height. The model predicted that *v*_H_ was positively related to *D* and irradiance, negatively to air temperature and CO_2_, and its negative correlations with *K*_S_ and ΔΨ_max_ (Fig. 1,S1). The sensitivities of maximum vapour pressure deficit (*D*_max_) and air temperature (*T*_max_), mean irradiance (*I*_abs_) during growing season to *v*_H_ variation after log-transformed were 0.6, 1 and –0.03 theoretically derived from our model, which implied that high *v*_H_ was expected at dry and cold areas with high irradiance. A detailed model description is presented in Methods, and derivation of the theoretical prediction is presented in Supplementary.

**Fig. 1.**
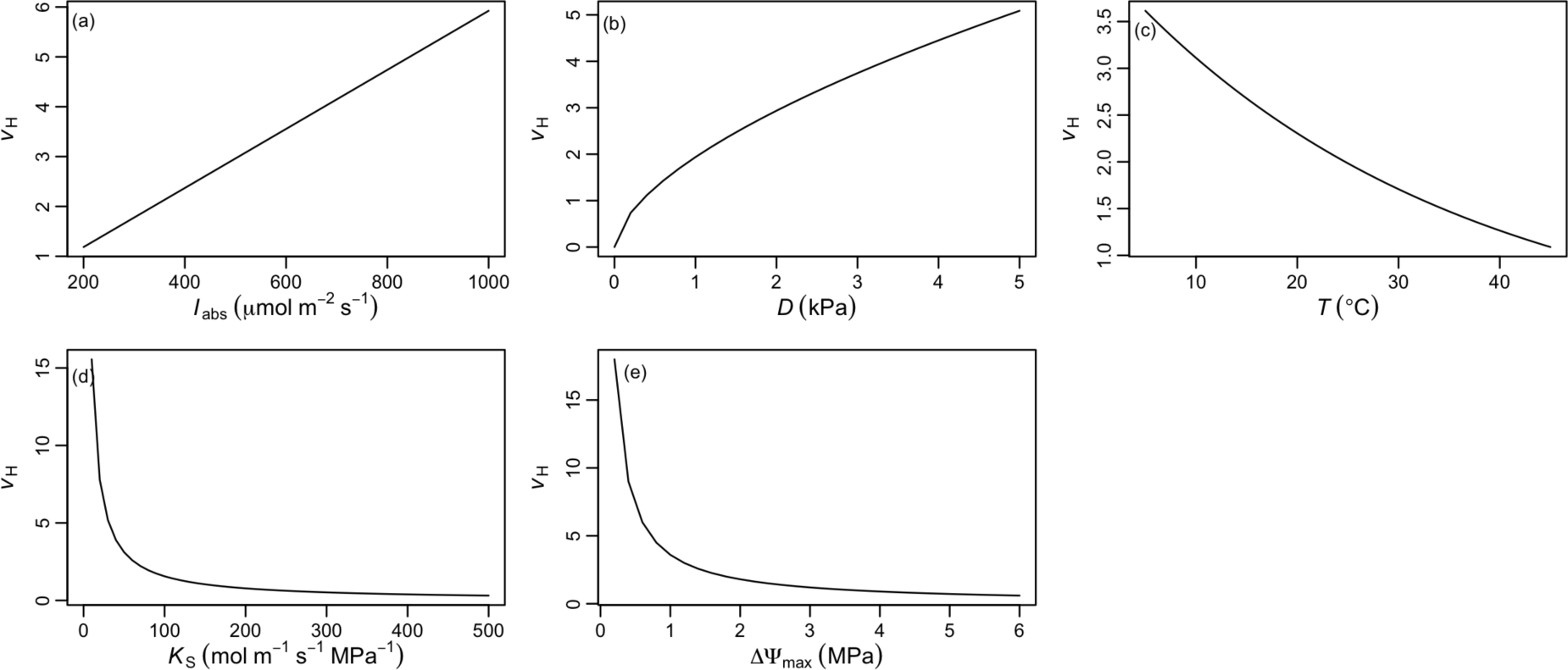
The sensitivities of theoretical model to climate and hydraulic traits. Sensitivity of the theoretical ratio of sapwood to leaf area (*v*_H_, 10^−4^) to irradiance (*I*_abs_, panel a), vapour pressure deficit (*D*, panel b), temperature (*T*, panel c), hydraulic conductivity (*K*_S_, panel d) and water potential difference between soil and leaf (ΔΨ_max_, panel e). Sensitivity analyses were done while keeping all other climate variables at median levels across species: *T* = 25.5 °C, *D* = 1.5 kPa.

The hypothesis thus for the first time explicitly incorporates the known functional coordination between hydraulic traits (i.e. the correlations between *v*_H_ and *K*_S_, ΔΨ_max_) in a parsimonious and analytical way, and quantitatively predicts the impacts of climates on this coordination network in terms of both the directions and sensitivities of those responses.

## Results

### The correlations between *v*_H_ and *K*_S_, ΔΨ_max_

We observed that greater *K*_S_ was associated with lower *v*_H_ and this trade-off was consistent across species and sites from the multiple linear regressions including climate variables (Fig. 2, 3). In multiple linear regressions including *K*_S_ and climate variables, the correlation between *v*_H_ and *K*_S_ was tighter at site level than species level. *K*_S_ was the most important driver of *v*_H_, explaining 46% of its variation at site level. The fitted *K*_S_ – *v*_H_ slopes (either at species or site level) were flatter than that predicted from theory (–1) (Fig. 2a, 3a). The fitted site-level slope was closer to theoretical prediction than that at species level. Analyses across a comprehensive sample of biomes and climate gradients (Fig. S2) showed that deciduous and evergreen species had similar mean value of *v*_H_ (2×10^−4^ and 1.6×10^−4^, respectively), but evergreen species had significantly lower *K*_S_ value than deciduous species (1.15 and 1.93 kg m^−1^ MPa^−1^ s^−1^, respectively, Fig. S3). In order to assess the variation in this trade-off among leaf phenology, the standard major axis (SMA) regressions were carried out for evergreen and deciduous species separately. However, the evergreen and deciduous species behaved the same and the slopes of their relationship were steeper than that from multiple linear regression as expected and not significantly different from the theoretical value (–1) (Fig. S5). Though few gymnosperms were included, the relationship between *v*_H_ and *K*_S_ was similar among angiosperms and gymnosperms (Fig. 3a). An insignificant relationship between *v*_H_ and ΔΨ_max_ was found at species and site level when ΔΨ_max_ was added into multiple linear regression (Fig. S6e), which might be masked by the strong negative relationship between *K*_S_ and ΔΨ_max_ (Fig. S7a). The relationship between *v*_H_ and ΔΨ_max_ could be indirect through the covariation with *K*_S_, which might be hard to examine empirically. The insignificant effect of ΔΨ_max_ may also attribute to the uncertainty in gridded soil water potential data (Fig. S7b).

**Fig. 2.**
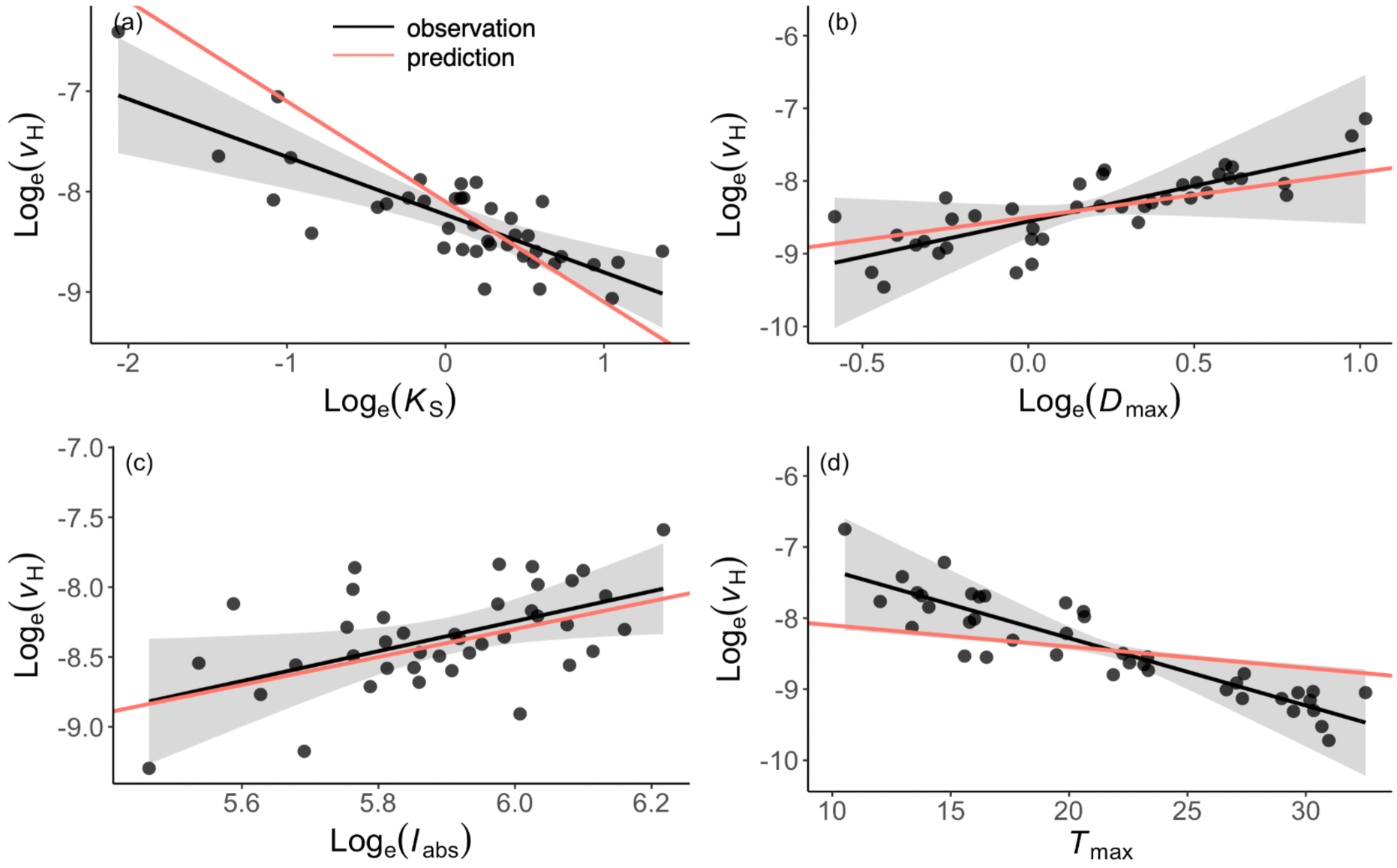
Partial residual plots from the multiple linear regression of log_e_-transformed the ratio of sapwood to leaf area (*v*_H_) against different predictors at site level using Dataset1. The predictors are shown in (a) sapwood-specific hydraulic conductivity (*K*_S_), (b) maximum vapour pressure deficit (*D*_max_), (c) mean irradiance (*I*_abs_), (d) maximum temperature (*T*_max_) during growing season. Black lines are the fitted across all sites and the gray shadings are the 95% confidence intervals around the fitted lines. The red lines are theoretical sensitivities in equation (1).

**Fig. 3.**
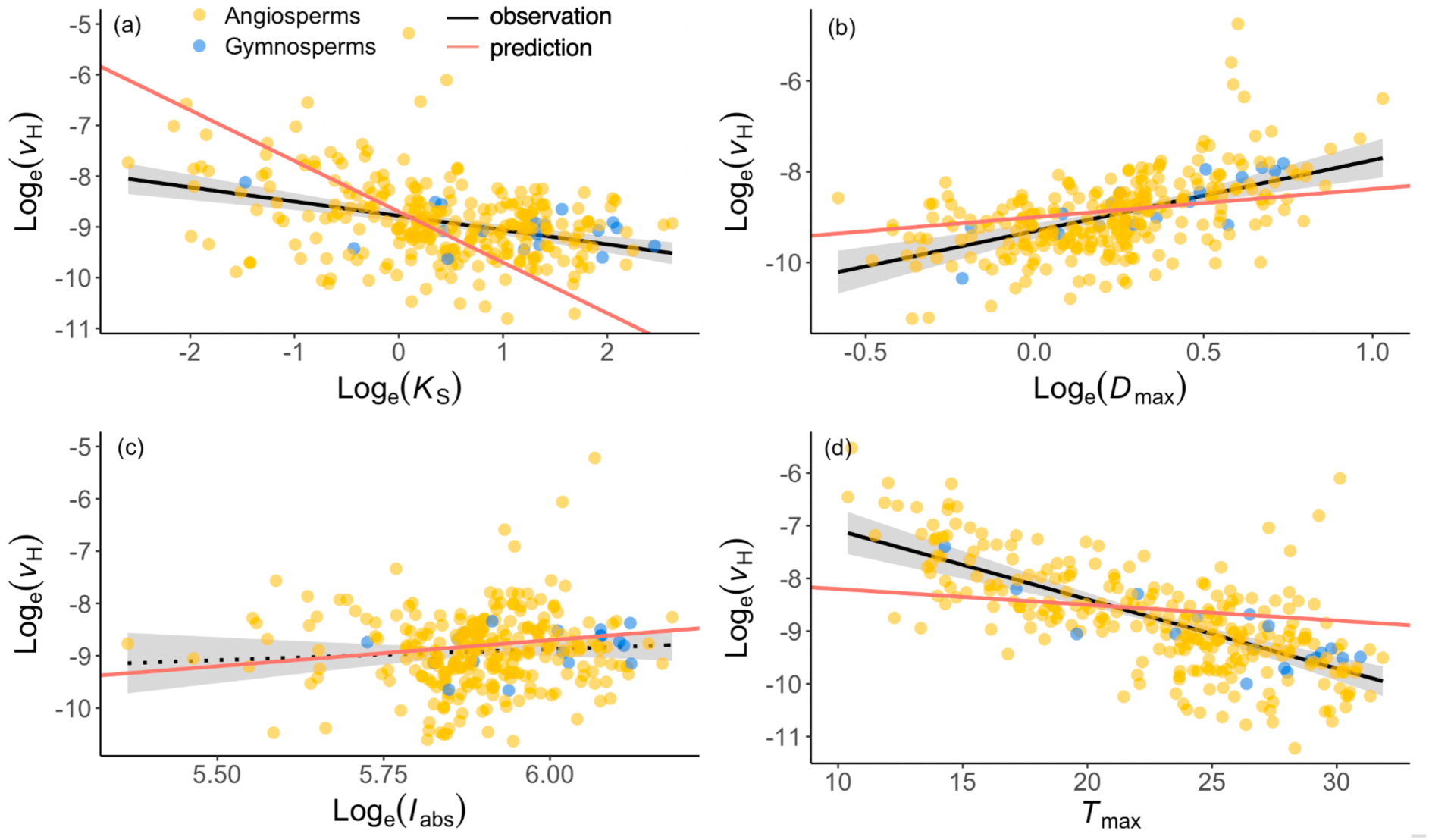
Partial residual plots from the multiple linear regression of log_e_-transformed the ratio of sapwood to leaf area (*v*_H_) against different predictors at species level using Dataset1. The predictors are shown in (a) sapwood-specific hydraulic conductivity (*K*_S_), (b) maximum vapour pressure deficit (*D*_max_), (c) mean irradiance (*I*_abs_), (d) maximum temperature (*T*_max_) during growing season. Black lines are the fitted across all sites and the gray shadings are the 95% confidence intervals around the fitted lines. The black solid lines are significant (*p*<0.05) and dotted line is insignificant (*p*>0.05). The red lines are theoretical sensitivities in equation (1).

### *v*_H_ variation along climate gradient

Across both species and site levels, we found that plants tended to have larger sapwood area and/or lower total leaf area as vapour pressure deficit and irradiance increased, and temperature dropped after consideration of traits correlation. The observed climatic effects on *v*_H_ variation were similar whether using species-averaged dataset (Dataset1) or individual-at-site dataset (Dataset2) (Fig. 3,S8). In Dataset1, the significant effects of temperature and *D*_max_ on *v*_H_ variation were stronger at site level than species level, but the effect of light was only significant at site level (Fig. 2,3,S6). The multiple linear regressions showed that these climatic effects together led to 23% more of *v*_H_ variation explained at site level (Fig. 3). The observed directions of climatic effects were consistent with EEO-based predictions and their predicted magnitudes fell into the confidence interval of slopes fitted at site level (Fig. 3b-d, S8b-d). The site-level observed effects of *D*_max_ and irradiance were close to theoretical sensitivities in species-averaged dataset (Dataset1, Fig. 3b,c). The observed effect of temperature in Dataset2 was smaller than that in species-averaged dataset and matched well with theoretical prediction (Fig. S8d).

### Predictability of hydraulic traits

The EEO-based model captured 56% of *v*_H_ variation using theoretical sensitivities of *K*_S_, irradiance, *D*_max_ and temperature, a fitted parameter (equation 12, Fig. 4a). The 46% of *v*_H_ variation was contributed from *K*_S_, followed by 6% from light, 4% from *D*_max_ and only 1% from temperature (Fig. S9). Substituting *K*_S_ with *v*_H_ in equation (12) could also lead to 66% of *K*_S_ variation predicted across sites (Fig. 4b).

**Fig. 4.**
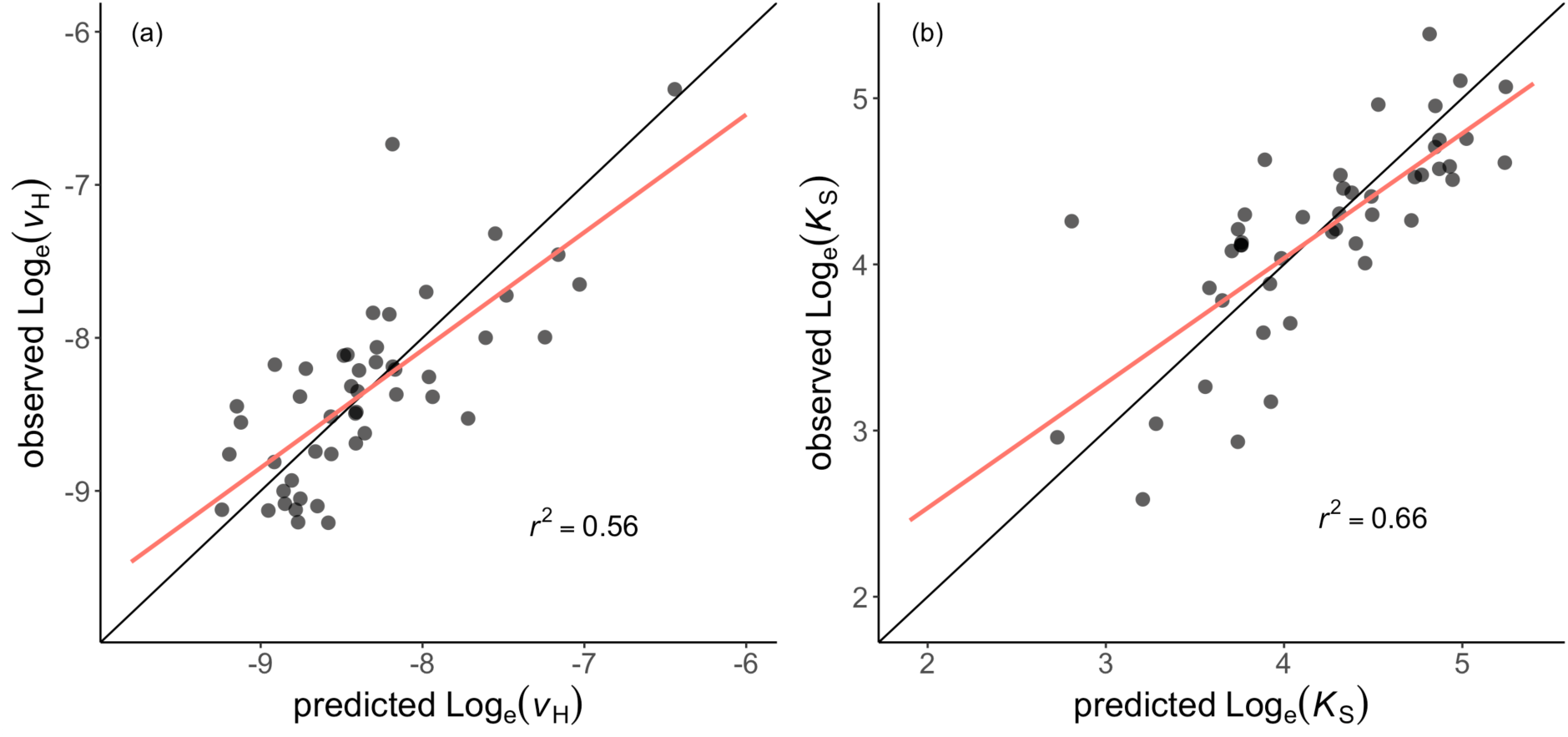
Comparison between site-mean observed and predicted ratios of sapwood to leaf area (*v*_H_) using Dataset1. (a) *v*_H_ is predicted using observed sapwood-specific hydraulic conductivity (*K*_S_) and climate variables with theoretical sensitivities of predictors and a fitted intercept. (b) *K*_S_ is predicted using observed *v*_H_ and the same climate variables.

## Discussion

The comparison between observation and prediction confirms the hypothesis that plants allocate resources optimally to balance the water supply through stem and water demand via leaf to maintain CO_2_ capture and photosynthesis without extra carbon wasted. We estimated the climate effects on *v*_H_ and global patterns of *v*_H_ for the first time by extending the model proposed by Whitehead et al. (1984) further and incorporating recently developed EEO-based models for photosynthetic traits. Our EEO-based theory, using a simple equation with only one fitted parameter to represent the coordination of hydraulic and photosynthetic processes, clearly describes the correlations between traits, quantifies the sensitivities of driving factors of *v*_H_ variation and largely captures its observed variation along climate gradients at a global scale. We show that hydraulic efficiency has a huge impact on *v*_H_ value globally and part of climate effects affects *v*_H_ indirectly through photosynthetic traits. The model also explains the smaller effect of temperature and precipitation on *v*_H_ when other traits are involved than that in a separate relationship solely including climate (Mencuccini et al. 2019). The change in *v*_H_ reflects the response of water balance within plant and carbon allocation to climate, which has great impact on the water and carbon cycling. This study helps explain how plants adjust to the changing climate via mediating resources and physiological processes, and emphasizing the importance of flexible trait representation to increase accuracy and reduce uncertainty under future scenarios in DGVMs and thus climate prediction in ESMs (Matheny et al. 2017; Scheiter et al. 2013; Berzaghi et al. 2020). The feature of simplicity allows rather easy incorporation of *v*_H_ without increasing model complexity.

*v*_H_ is directly associated with plant transpiration and regulates water cycling from soil to atmosphere. The sensitivity of *v*_H_ to climate is important for us to understand how water cycling at plant level reacts to the climate change. Our estimated sensitivities help us elucidate the interannual response of plant water cycling under climate change. In addition to the widely acknowledged positive effect of vapour pressure deficit (*D*), our theory predicts a negative temperature and positive irradiance effects on *v*_H_, which is consistent with the observation that plants with higher *v*_H_ value are observed at dry and cold area with high light availability using two dataset at species and individual levels (Fig. 2, 3, S9). Globally increasing vapour pressure deficit (*D*) poses a dominant impact on leaf water potential regulated by stomata to control the water demand or leaf gas exchange (Turner et al. 1984; Yuan et al. 2019; Fang et al. 2022). Therefore, *D* influences whole plant hydraulic streams through the gradient from soil to leaf. In addition, the induced changes in stomatal conductance alter CO_2_ uptake and photosynthetic process, leading to the shift in water demand and *v*_H_. Other studies also find higher *v*_H_ value at sites with low soil water availability (Rosas et al. 2019). On the one hand, soil water availability is related to soil water potential and changes ΔΨ_max_. This may further influence carbon allocation between belowground and aboveground (more carbon invested in roots when soil water availability is low). On the other hand, it covaries with atmospheric dryness and hard to disentangle their effects (Fu et al. 2022).

The collinearity of *D* and temperature also makes it difficult to separate their individual effect with even divergent results from warming experiments (Way et al. 2013; Mcbranch et al. 2018). The EEO-based model allows to disentangle their effects theoretically and reveals the opposite impacts of *D* and temperature on *v*_H_, which emphasizes the importance of quantifying their sensitivities to predict plant allocation strategy under future climate when temperature and *D* both increase (Fig. S10). The total temperature effect is the sum of its effect on photosynthetic trait and hydraulic efficiency (see details in SI). The response of photosynthetic traits to temperature has been widely studied. As temperature increases, photosynthetic capacity increases and the ratio of leaf internal to external CO_2_ becomes larger, leading to higher stomatal conductance and water demand. At the same time, hydraulic efficiency increases due to low water viscosity and high permeability in symplastic pathway that is rarely studied and poorly understood. Here we employed a rather conservative temperature dependency for hydraulic efficiency (Matzner et al. 2001; Cochard et al. 2000). This indicates that theoretical temperature sensitivity on *v*_H_ may be greater if less conservative temperature sensitivity was applied. Our results highlight the need to study the temperature response of hydraulic efficiency in more detail, due to its important role in constraining photosynthesis process, especially under global warming.

The impact of irradiance is often overlooked in previous analysis solely about hydraulic traits, however hydraulic processes are tightly coupled with photosynthetic process, leading to the crucial role of light in influencing hydraulic traits. The observation and prediction from our EEO-based model both find plants need more sapwood area to support the same leaf area when irradiation is high. More water is required to maintain high assimilation rate under high irradiation in order to utilize light optimally at the whole-plant level. Plants may have greater leaf vein length per unit area to match increased water supply through xylem (Sack et al. 2013). The integration of hydraulic and photosynthetic processes can help us better comprehend plant strategy from a more unified perspective. Our EEO-based theory predicts that plants support more leaves without allocating extra carbon to sapwood under higher CO_2_ by increasing water use efficiency. We do not examine the CO_2_ effect here due to limited hydraulic data along a CO_2_ gradient, however our predictions match the direction of trend found in other studies (Westoby et al. 2012; Trugman et al. 2021).

Our hypothesis for *v*_H_ variation is tested temporally at species and site level, indicating that our model works well on a longer timescale (multiple years) that involves some evolutionary processes, such as plants acclimation and species turnover. Theoretically, photosynthetic traits in our model acclimate to climate on a weekly timescale, while the acclimation of hydraulic traits is poorly known. The shortest time frame for *v*_H_ to reach its optimal value to reconcile different timescales from water supply and demand still needs further verification. Previous studies show that plants can increase *v*_H_ to cope with drought event by shedding leaves to prevent water loss or hydraulic failure during a rather short time period (Carnicer et al. 2011; Choat et al. 2018; Trugman et al. 2018). During drought, photosynthetic traits can be adjusted quickly to alter water demand (Mengoli et al. 2022), but maximum hydraulic conductivity is not achieved and water storage within stem may be released, leading to disequilibrium between water demand and supply. A recent photosynthesis model coupled with plant hydraulics adopts the hypothesis that water supply matches water demand at any time, which well predicts photosynthetic processes under drought on a daily timescale (Joshi et al. 2022). This indicates that our model shows the potential to perform at a short timescale and be extended to improve the representation of phenology in DGVMs (Martín Belda et al. 2022), which still requires more studies and tests on plant water transport and our model under stressed conditions on a weekly timescale.

Current DGVMs adopt fixed or flexible carbon allocation to leaf and stem mostly based on plant functional type parameters or empirical relationships in response to environment (Trugman et al. 2019; Berzaghi et al. 2020), leading to the uncertainty of land carbon sink (Sitch et al., 2008, O’Sullivan et al. 2022). *v*_H_ represents carbon allocation between leaf and stem, which is shown to vary with climate. The allocation to leaf and stem scheme directly influences the amount of low-turnover carbon stored in stem and quick-turnover of leaf for productivity and transpiration, which in turn, affects land carbon and water fluxes. Under future climate warming scenario, increasing CO_2_ and temperature can alleviate part of positive effect of *D* on *v*_H_, resulting in more leaves supported for a given stem. The change rate in climate variables along with their sensitivities of *v*_H_ together determine the carbon allocation in the future. Low *v*_H_ may lead to more CO_2_ captured from the atmosphere and slow down climate warming. Our EEO-based model for *v*_H_ provides a new route of flexible carbon allocation scheme in DGVMs/ESMs to generate more realistic model output (Magnoni et al. 2000; Deckmyn et al. 2006; Trugman et al. 2019).

With increasing *D* and drought events, the importance of hydraulic traits for improving the prediction of vegetation productivity and transpiration has been recognized. With more vegetation models incorporating additional plant hydraulic processes, maximum plant hydraulic conductivity becomes an important model input for transpiration and carbon assimilation calculation. The lack of mechanism of hydraulic trait variation hinders the model performance at a large scale or leads to fixed hydraulic trait value for PFTs, which increases model uncertainty (Christoffersen et al. 2016; Matheny et al. 2017; Venturas et al. 2018; Sperry et al. 2017; Kennedy et al. 2019; Eller et al. 2020). Due to the time-consuming measurement of hydraulic conductivity, its availability is limited globally (Kattge et al. 2020). Whereas, *v*_H_ is easier and quicker to measure in the field. In a trait-driven model, the empirical relationship is applied to simulate hydraulic conductivity using *v*_H_ (Christoffersen et al. 2016). In our study, the theory derives the simple relationship between *v*_H_ and *K*_S_, which shows the successful prediction of *K*_S_. This demonstrates the potential to substitute the hydraulic PFTs in DGVMs and ESMs to perform at a larger scale. The inclusion of *K*_S_ variation along climate gradient in DGVMs changes the water supply that constrains gas exchange, leading to accurate transpiration and carbon assimilation at regional scale.

This robust trade-off between *v*_H_ and *K*_S_ across site and species has long been recognized as compensation effect, indicating its essential role in maintaining plant fitness. It is consistent with the prediction of our model that the relationship between *v*_H_ and *K*_S_ should be negative and modified by climate and other traits. Hydraulic efficiency is fundamental for *v*_H_ due to its direct constrains on water supply within plants. When more leaves are produced, plants either increase conducting area or efficiency to maintain water supply when assimilation rate and stomatal control remain constant. This compensation effect guarantees the balance between water supply though xylem and water demand from leaf to avoid wasted carbon investment or protect xylem from excessive tension. The importance of this compensation effect also manifests in their evolutionary correlation at species level, which means *v*_H_ and *K*_S_ cooperatively adapt to the changing environment over long timescale (Sanchez-Martinez et al. 2020). This implies that the trade-off between *v*_H_ and *K*_S_ in our model still persists under future climate conditions. Such key trade-off could be used to constrain species properties in individual-based vegetation models (Berzaghi et al. 2020).

Our EEO-based theory expects a negative relationship between *v*_H_ and ΔΨ_max_, nonetheless, this correlation might be partially concealed by the significant relationship between ΔΨ_max_ and *K*_S_ at the global dataset. The effect of ΔΨ_max_ is the combination of soil water potential (Ψ_soil_) and leaf water potential at turgor loss point (Ψ_tlp_), which is directly related to stomatal behaviour. Leaf turgor has shown to be the dominant contributor to changes in stomatal conductance, and Ψ_tlp_ well represents the point when stomata fully close and carbon assimilation stops (Cochard et al. 2002; Bartlett et al. 2016; Rodriguez-Dominguez et al. 2016; Knipfer et al. 2020). Soil water potential influences the water potential from soil to stem to regulate stomatal conductance (Rodriguez-Dominguez and Brodribb 2020). Plants maintain the water transport through water potential gradient between soil and leaf, avoid excessive water loss and xylem dysfunction via stomata regulated by leaf turgor. However, different stomatal conductance models show large variation in its response to soil water potential (Sabot et al. 2022). Some DGVMs including plant hydraulics employ another important hydraulic trait (Ψ_50_, water potential at 50% loss of hydraulic conductivity) to represent the species’ response to drought via its relationship with hydraulic conductivity (Christoffersen et al. 2016; Sperry et al. 2017; Eller et al. 2018). Whereas, Brodribb et al. (2003) finds no correlation between Ψ_50_ and stomatal closure point across species. Previous studies demonstrate stomata closure occurs before Ψ_50_ or xylem cavitation (Bartlett et al. 2016; Martin-StPaul et al. 2017). Although the models demonstrate the potential to capture plants response under drought, it neglects the mechanism that stomata are closely regulated by ΔΨ_max_ rather than hydraulic conductivity. The direct incorporation of Ψ_tlp_ effect into this model increases its realism and robustness of stomatal behaviour under drought event, but we currently excluded it when *v*_H_ was predicted due to the large uncertainty in Ψ_soil_ (Fig. S) extracted from gridded product (Hengl et al. 2017).

Compared to other models, our EEO-based model is validated spatially at longer timescale and considers coordinated physiological processes to achieve an easy equation with single parameter. Although the method of maximizing net carbon gain over optimal *v*_H_ could explain its response to *D* and CO_2_ effect, the model may also choose other traits or processes to maximize, such as commonly used stomatal behaviour (Sperry et al. 2017; Wang et al. 2020), which may lead to changes in *v*_H_. Whereas, our model is based on the hypothesis that the input should match output of a system to maintain its function and stability, this idea has been applied in other fields, such as Ohm’s law. Our model avoids the dependency of complicated processes that requires simplifications and parameters, which is beneficial for improvement of PFTs scheme in DGVMs without jeopardizing original model structure and increasing process complexity. Inclusion of such a parsimonious scheme in DGVMs and ESMs will lead to improved representation of plant responses to future climate change, with projected increases in temperature, VPD, regional drought, and extreme events, and thus in turn a critical for improved future climate prediction.

## Method

### Theory based on eco-evolutionary optimality

The model of *v*_H_ variation builds on the hypothesis proposed by Whitehead et al. (1984) and Xu et al. (2021), which predicts that maximum plant water transport should match maximum photosynthesis with resources optimally allocated. This implies that water loss through stomata should match water transport through xylem, which can be estimated using Fick’s and Darcy’s law respectively. The use of water storage and xylem water refilling are not considered in our model as they often occur during abrupt extreme events and night.

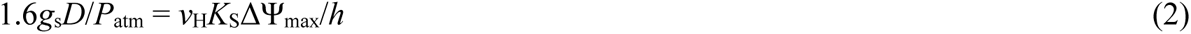

where *g*_s_ is stomatal conductance to CO_2_ (mol m^−2^ s^−1^), *D* is the vapour pressure deficit (Pa) and *P*_atm_ is the atmospheric pressure (Pa). On the right-hand side of the equation, *v*_H_ is the ratio of sapwood to leaf area (m^2^ m^−2^); *K*_S_ is the maximum sapwood-specific hydraulic conductivity (mol m^−1^ s^−1^ MPa^−1^); ΔΨ_max_ is the maximum difference between leaf and soil water potential (Ψ_min_ and Ψ_soil_, MPa); *h* is the path length, approximately equal to tree height (m).

We can calculate *g*_s_ from the diffusion equation and the photosynthesis model (Farquhar et al., 1980):

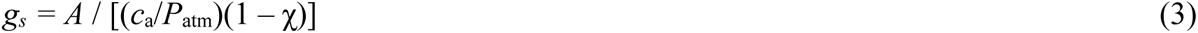

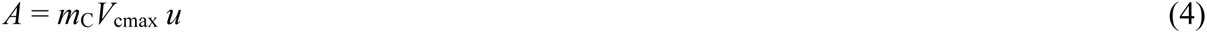

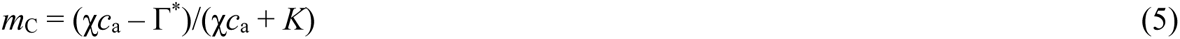

where *A* is the assimilation (photosynthesis) rate (mol m^−2^ s^−1^), *c*_a_ is the ambient partial pressure of CO_2_ (Pa), χ is the ratio of leaf-internal to ambient CO_2_ partial pressure (Pa Pa^−1^), *V*_cmax_ is the maximum capacity of carboxylation (mol m^−2^ s^−1^), Γ* is the photorespiratory compensation point (Pa), and *K* is the effective Michaelis-Menten coefficient of Rubisco (Pa), *u* is the unit conversion from μmol to mol. By replacing *g*_s_ from equations (3)-(5) in equation (2) and rearranging after Log_e_ transformation, we derive the following equation (6) that describes the coordination between hydraulic and photosynthetic traits:

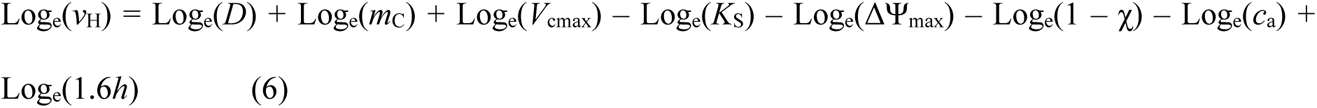

We can estimate photosynthetic traits (χ, *V*_cmax_) using existing models based on EEO. The least-cost hypothesis states that plants minimize the unit cost of both photosynthesis and transpiration (Prentice et al. 2014; Wang et al. 2017), leading to the prediction of χ (equation 7).

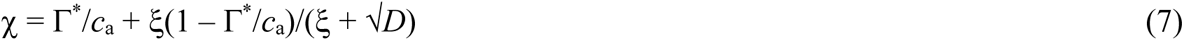

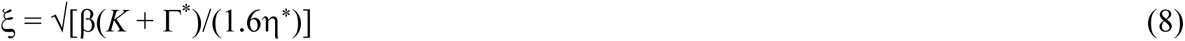

where β is a dimensionless constant (146, based on a global compilation of leaf δ^13^C measurements), and η* is the viscosity of water relative to its value at 25 °C.

The coordination hypothesis states that light- and Rubisco-limited photosynthesis rates should match to optimally utilize light without extra maintenance costs (Smith et al. 2019), leading to the prediction of *V*_cmax_ (equation 9).

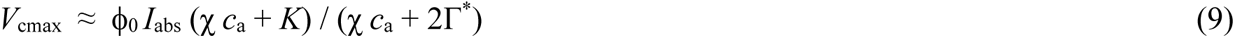

where ϕ_0_ (μmol C μmol^−1^ photon) is the intrinsic quantum efficiency of photosynthesis calculated by temperature-dependence relationship (Bernacchi et al. 2003), *I*_abs_ is the photosynthetic photon flux density absorbed by leaves (μmol m^−2^ s^−1^).

Since photosynthetic traits can be estimated from climate variables alone, the relationship between *v*_H_ and its drivers can be represented in equation (10). Thus, the sensitivities of climate variables can be derived from the photosynthesis optimality models (see detailed steps in supplementary), which results in the simple form of our theoretical model for *v*_H_ (equation 1).

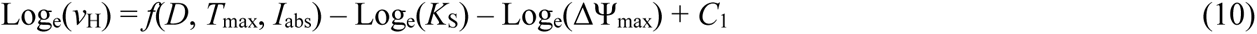

### Datasets

The global species-averaged dataset (Dataset1) for 1727 species was an updated version of HydraTRY (Mencuccini et al. 2019; Sanchez-Martinez et al. 2020; See Supplementary). It includes 1624 angiosperms and 103 gymnosperms, 332 deciduous and 694 evergreen species with remaining species unknown. Species names and taxonomy (angiosperm and gymnosperm) were checked against the Plant List using *plantlist* package. The information of leaf habit was matched with original publications and online reports. The dataset contained the branch-based ratio of sapwood to leaf area (*v*_H_, m^2^ m^−2^, dimensionless), maximum sapwood-specific hydraulic conductivity (*K*_S_, mol m^−1^ s^−1^ MPa^−1^) and Ψ_tlp_ (MPa). The traits values were averaged if multiple samples for one species occurred in order to achieve maximum number of species with available all traits values at the same time.

Monthly climate data for Dataset1 from year 1970 to 2000 was extracted from Worldclim interpolated from weather stations (Fick and Hijmans, 2017), including maximum temperature, mean temperature, vapour pressure, solar radiation and precipitation. Monthly maximum vapour pressure deficit (*D*_max_) was calculated using maximum temperature and vapour pressure. Monthly volumetric soil water content (m^3^ m^-3^) of the ECMWF Integrated Forecasting System for 7-100cm depth from 1981-2019 (https://cds.climate.copernicus.eu/#!/home), soil texture, soil organic carbon content were extracted from SoilGrids (https://soilgrids.org/) to generate soil water potential (Ψ_soil_) using *medfate* R package (Saxton and Rawls, 2006). Monthly aridity index (AI, the ratio of precipitation to evapotranspiration) was obtained from Global Aridity and PET Database (https://cgiarcsi.community/data/global-aridity-and-pet-database/) to determine growing season for plants. We defined the growing season as the months when mean temperature was above 0°C and aridity index was above 0.1 and limited our study data to these periods, as in temperate ecosystems at periods below these values our study species will either be leafless or not photosynthesising. Maximum temperature (*T*_max_), vapour pressure deficit (*D*_max_), mean photosynthetically active radiation (*I*_abs_) and Ψ_soil_ during growing season were calculated. The maximum difference between leaf and soil water potential (ΔΨ_max_) was estimated using Ψ_tlp_ and soil water potential during growing season. The climate data for each species were calculated as the mean value of the per pixel value across its spatial extent within the observational data.

The global dataset (Dataset2) of 1612 individual samples (1247 species) was compiled from published literatures for validation of our results without aggregating into species level (See Supplementary). Most of *K*_S_ data came from He et al. (2020) and corresponding *v*_H_ values for each record have been implemented from the original publications. We added more *K*_S_ and *v*_H_ data from 2017-2021 by conducting searches on Web of Science, Google Scholar, and China National Knowledge Infrastructure (http://www.cnki.net) using the keywords “leaf area to sapwood area ratio”, “Huber value”, and “hydraulic traits”. To minimize the errors from ontogenesis and methodology, we excluded the data that failed to meet the following criteria: (a) wild plants growing in natural ecosystems without experiments; (b) measurements were made on adult plants or saplings; (c) *v*_H_ measured on terminal stem or branch segments at top canopy; (d) *v*_H_ was estimated as the mean value for each species at the same site when the data was available for more than one individual. Because the corresponding Ψ_tlp_ data was limited, we did not include it in Dataset2. Monthly climate data from year 2011 to 2020 was extracted from CRU (https://crudata.uea.ac.uk/cru/data/hrg/, Harris et al. 2020) for each site, including maximum temperature, vapour pressure, cloud cover and precipitation. We calculated growing-season *T*_max_, *D*_max_ and *I*_abs_ using Simple Process-Led Algorithms for Simulating Habitats (SPLASH) model (Davis et al. 2017).

### Statistical analysis

We used Dataset1 at species level to test our EEO-based hypothesis by investigating how *v*_H_ changed along environmental gradient and comparing with model predictions. Principal components analysis (PCA) was conducted to analyse climate covariance and reduce three climate variables (temperature, irradiance and *D*) to two axes. The first axis of Principal Component Analysis (PCA) explained 74.6% of climate variation, which was dominated by maximum vapour pressure deficit (*D*_max_) and temperature (*T*_max_) during growing season (Fig. S4). The second axis (accounting for 20% of variation) largely reflected variation in site irradiance during growing season (*I*_abs_). In order to find the clear pattern of *v*_H_ variation in climate space, species with available three hydraulic traits values were divided into 48 climate zones/sites according to the loadings of first and second PC axes to investigate trait variation at site level (Fig. S4a). The relationships between *v*_H_ and its driving factors were analysed at species and site levels.

To examine the relationships between *v*_H_ and its driving factors in a linear format, the multiple linear regression was carried between *v*_H_ and *K*_S_, *T*_max_, *D*_max_, *I*_abs_ and ΔΨ_max_ at species and site level using Dataset1 after trait values were natural log-transformed. Due to data absence of one or more trait for some species, sites with only one species were excluded from the multiple linear regression. The regression was repeated without ΔΨ_max_ due to its insignificant effect on *v*_H_ variation and large uncertainty. Due to the unknown information about path length for water transport, the intercept (*C*_2_ = –9.59) in equation (12) was fitted for *v*_H_ prediction at site level when the coefficients of other factors were kept as theoretical values. *C*_2_ was the only parameter in our EEO-based model and contained the information about path length, photosynthetic traits values at standard conditions and unit conversion. The sensitivity plots of *v*_H_ to its driving factors were drawn when one factor changed at a time while others were kept as median of their observed values. To quantify the contribution of different factors in *v*_H_ prediction, the *r*^2^ of observed and predicted *v*_H_ with one factor added each time was calculated.

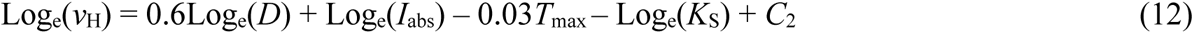

The multiple linear regression between *v*_H_ and *K*_S_, *T*_max_, *D*_max_, *I*_abs_ was conducted repeatedly using Dataset2 at site level for validation.

To assess the trade-off between *v*_H_ and *K*_S_ among different leaf phenologies (evergreen and deciduous), their bivariate relationship was examined using standardised major axis (SMA) regression and the slopes were tested if different from theoretical values (–1) or not using *smatr* package. To show the opposite effects of temperature and vapour pressure deficit on *v*_H_ variation clearly, we estimated *v*_H_ value in continuous climate space (temperature and vapour pressure deficit) using theoretical sensitivities of *D* temperature, while kept *K*_S_ and *I*_abs_ as median values.

## Author contribution

H.X., H.W., I.C.P., S.P.H., L.R., and M.M. designed the study. H.X., H.W. and I.C.P. developed the theory. M.M. and P.S. compiled hydraulic trait dataset and extracted climate data (Dataset1). P.H. and Q.Y. compiled hydraulic trait dataset (Dataset2). H.X. performed the analysis and wrote the first draft of the manuscript; all authors contributed to the final draft.

**Fig. S1.**
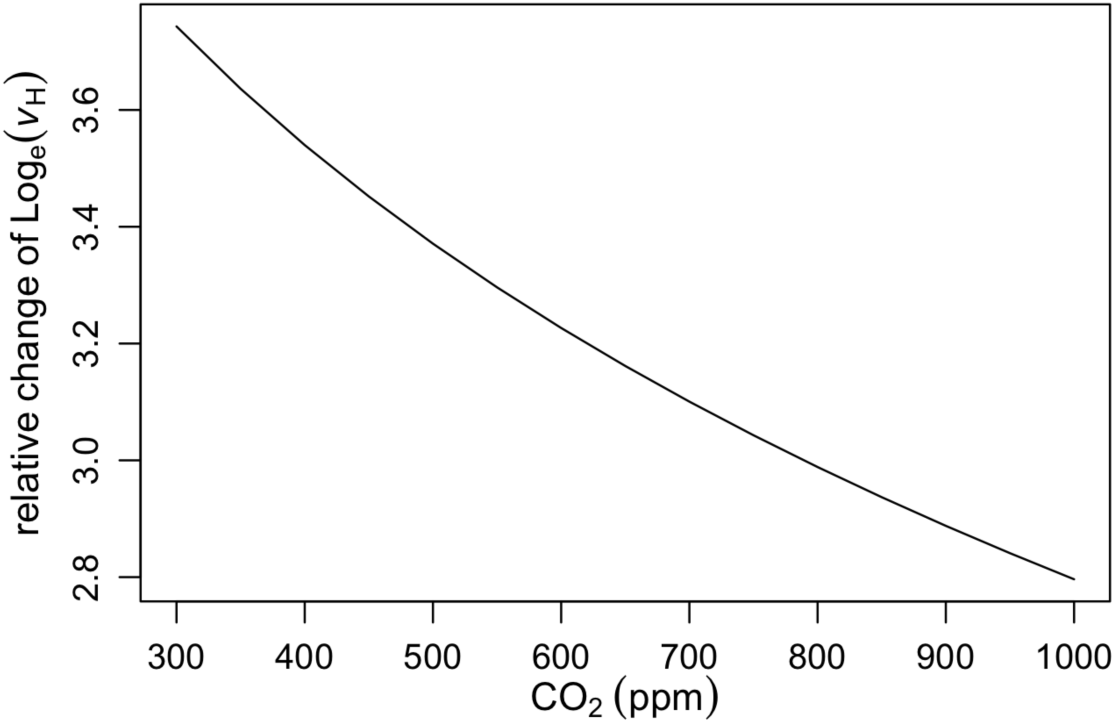
The theoretical sensitivity of log_e_-transformed ratio of sapwood to leaf area (*v*_H_) to ambient CO_2_.

**Fig. S2.**
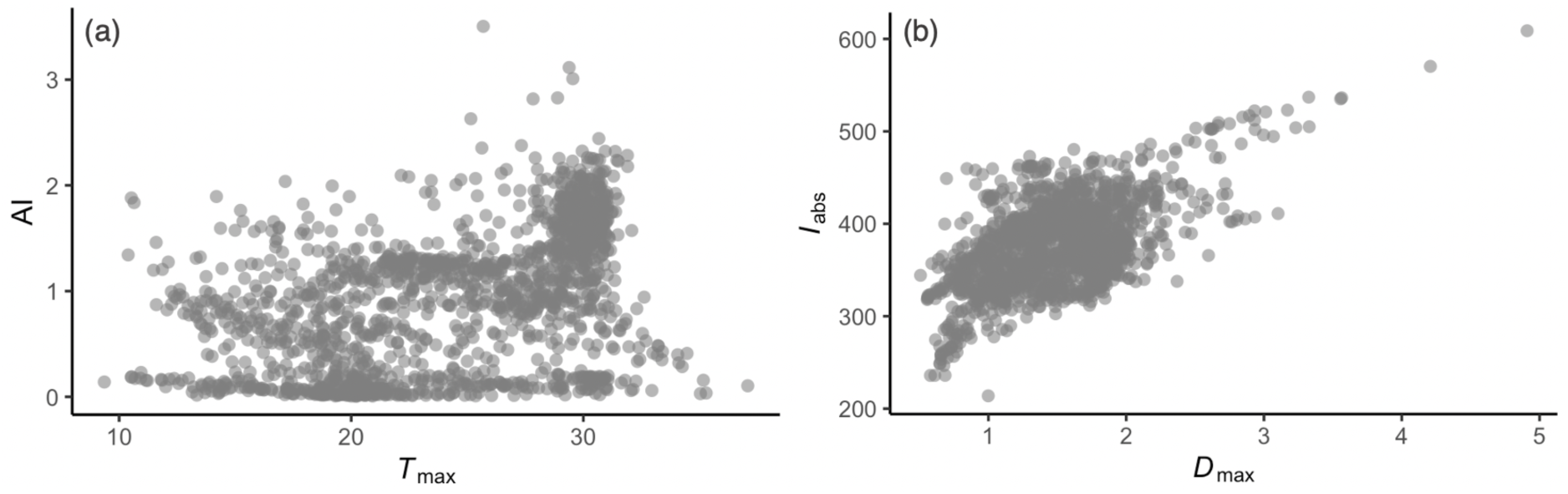
Species distribution along climate gradients in Dataset1. AI is aridity index (ratio of precipitation of evapotranspiration), *T*_max_ is maximum temperature, *D*_max_ is maximum vapour pressure deficit and mean irradiance (*I*_abs_) during growing season.

**Fig. S3.**
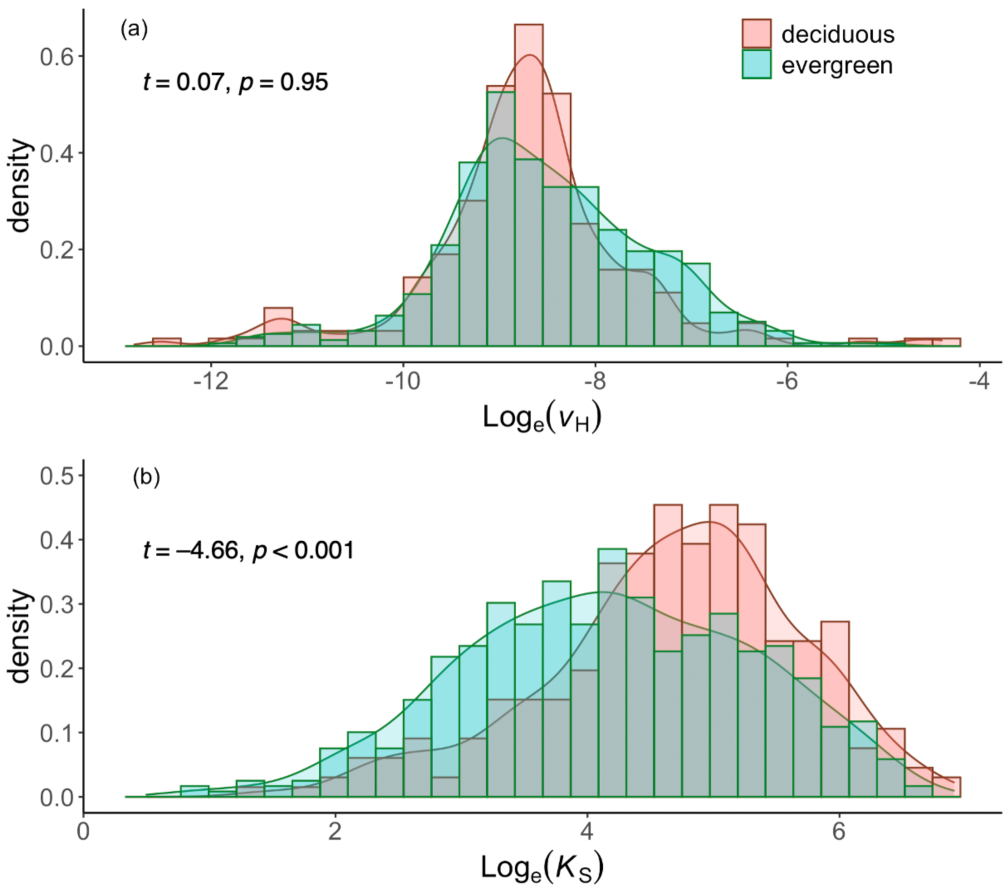
The distribution of ratio of sapwood to leaf area (*v*_H_) and hydraulic conductivity (*K*_S_) among deciduous and evergreen species in Dataset1. T-test is carried out to examine the difference of *v*_H_ and *K*_S_ between leaf phenology.

**Fig. S4.**
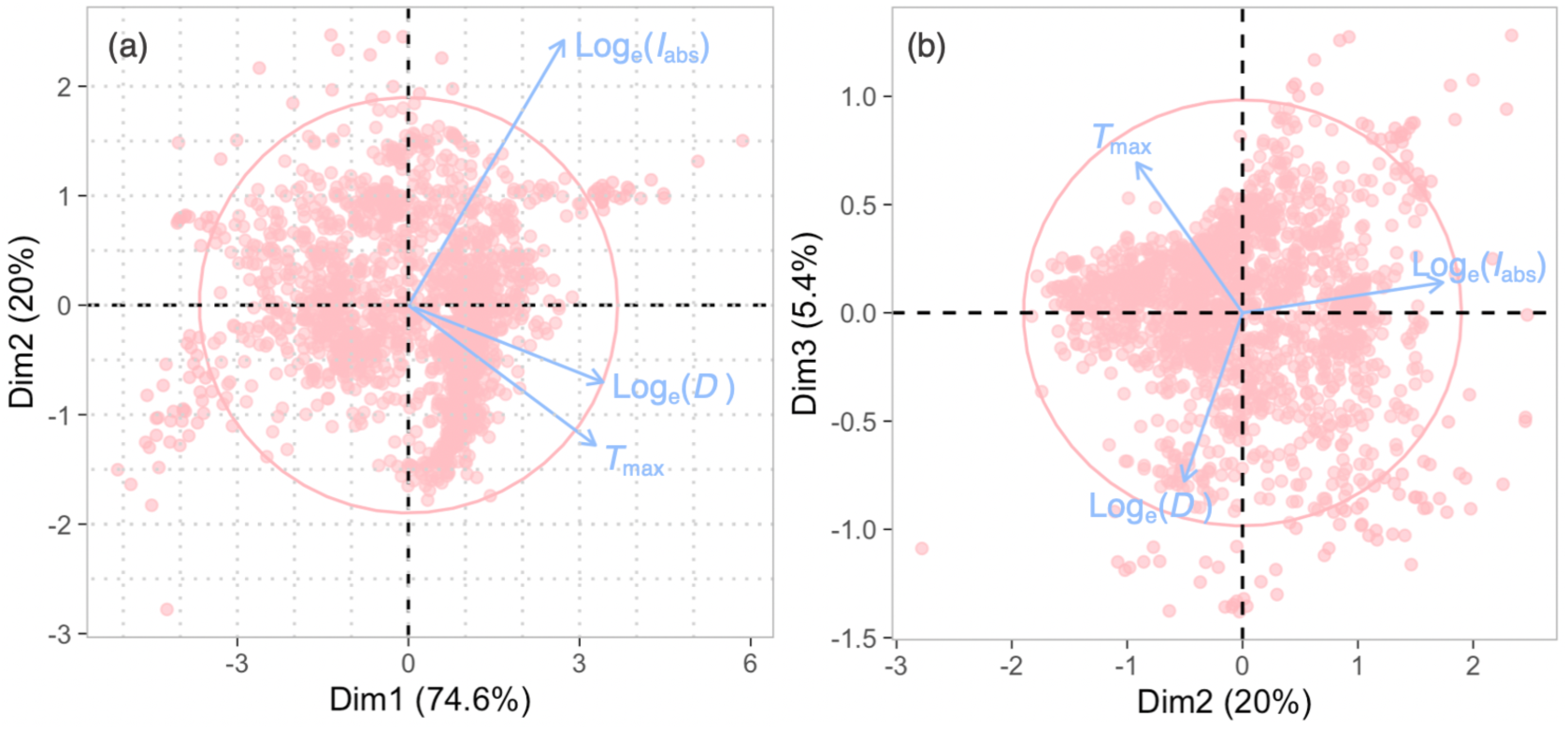
Principal components analysis of climate variables including maximum vapour pressure deficit (*D*_max_), maximum temperature (*T*_max_) and mean irradiance during growing season in Dataset1. (a) PC1 vs PC2, gray dotted lines show the site division according to loadings of PC1 and PC2 axes. (b) PC2 cs PC3. The pink points represent species.

**Fig. S5.**
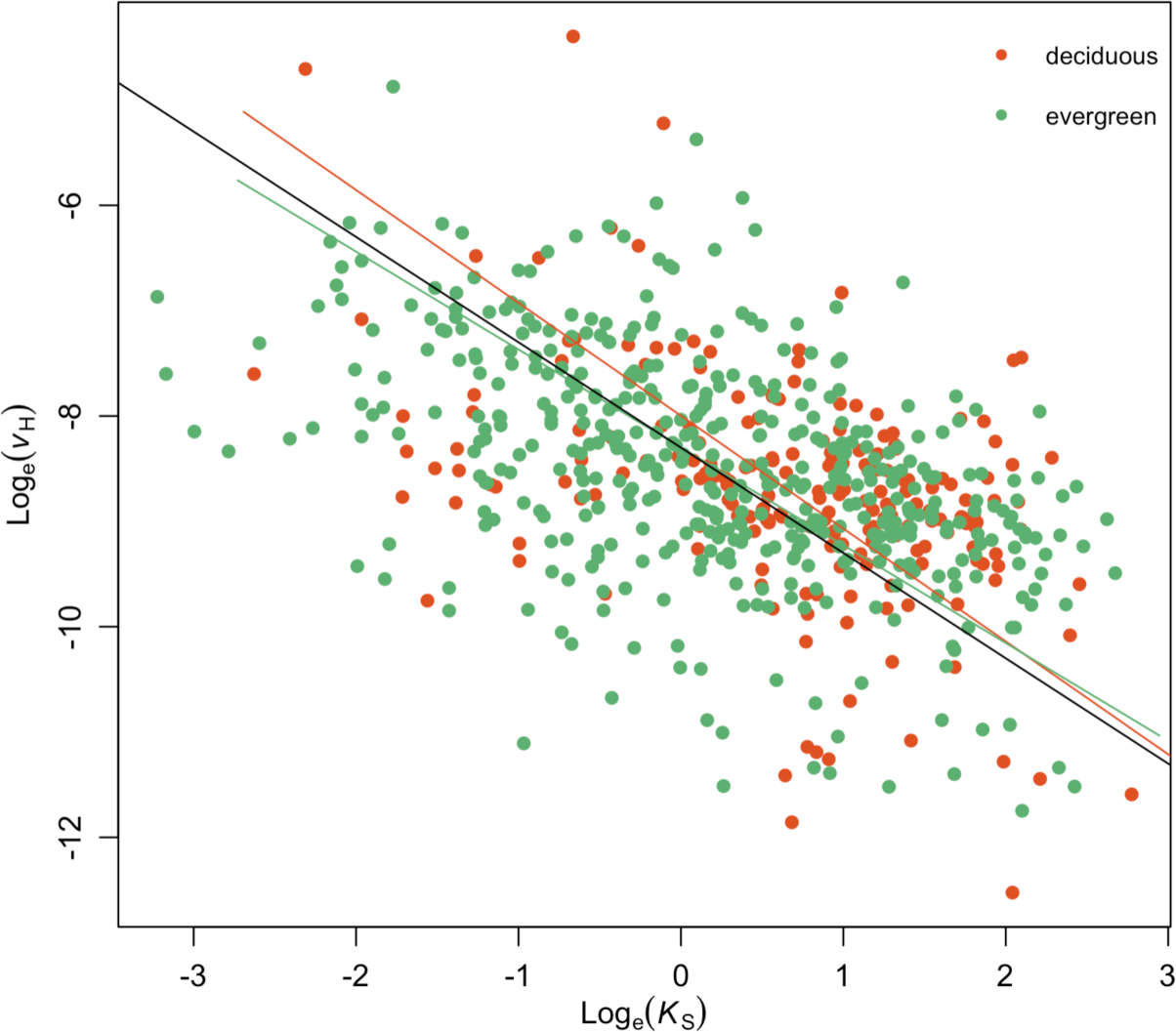
The relationships between log_e_-transformed ratio of sapwood to leaf area (*v*_H_) and hydraulic conductivity (*K*_S_) at species level using Dataset1. The red dots are deciduous species, green dots are evergreen species. The black line is 1:1 line, the red and green lines are fitted using standardised major axis (SMA) regression among deciduous and evergreen species respectively.

**Fig. S6.**
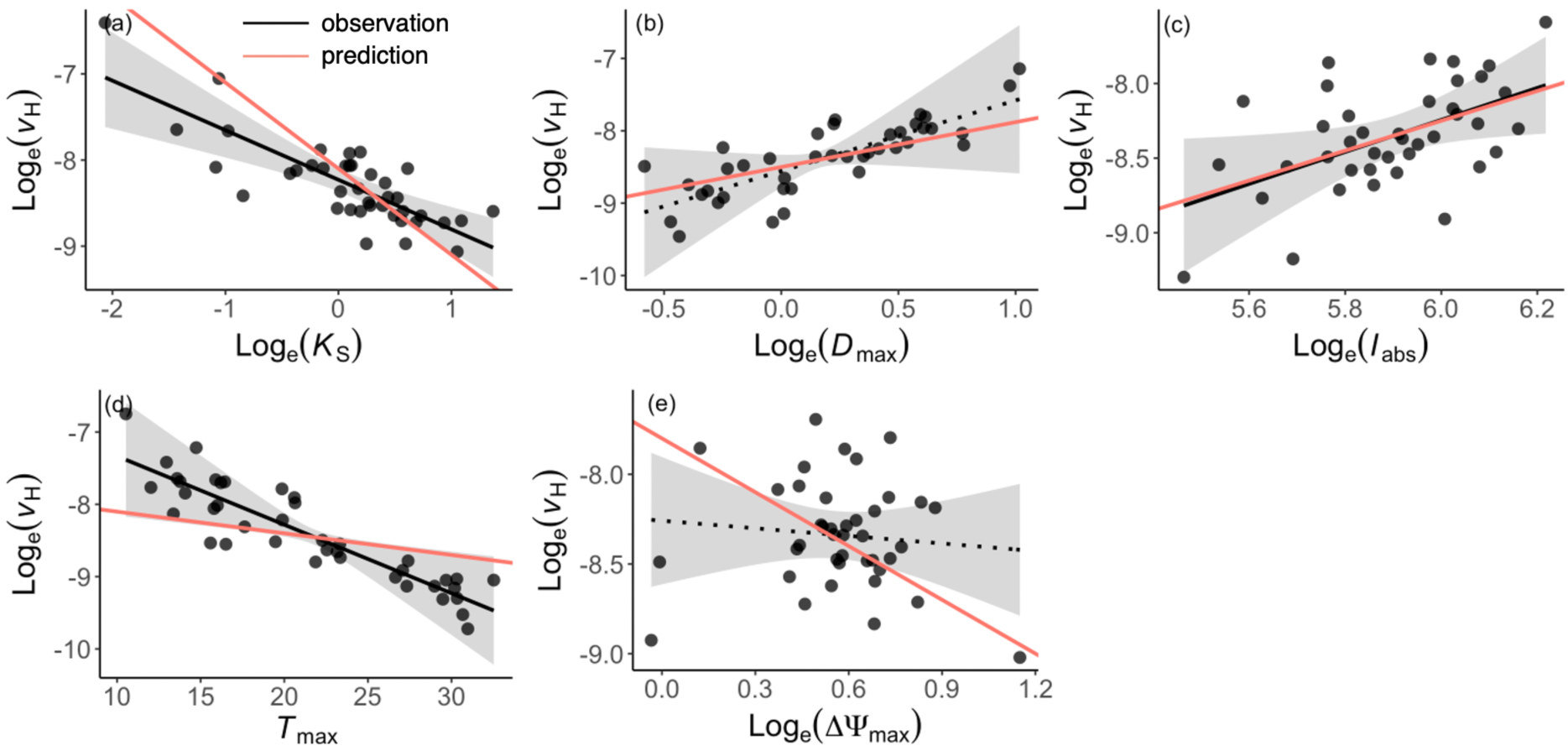
Partial residual plots from the multiple linear regression of log_e_-transformed the ratio of sapwood to leaf area (*v*_H_) against different predictors at site level using Dataset1. The predictors are shown in (a) sapwood-specific hydraulic conductivity (*K*_S_), (b) maximum vapour pressure deficit (*D*_max_), (c) mean irradiance (*I*_abs_), (d) maximum temperature (*T*_max_) and (e) maximum water potential difference between soil and leaf (ΔΨ_max_) during growing season. Black lines are the fitted across all sites and the gray shadings are the 95% confidence intervals around the fitted lines. The black solid lines are significant (*p*<0.05) and dotted line is insignificant (*p*>0.05). The red lines are theoretical sensitivities in equation (1).

**Fig. S7.**
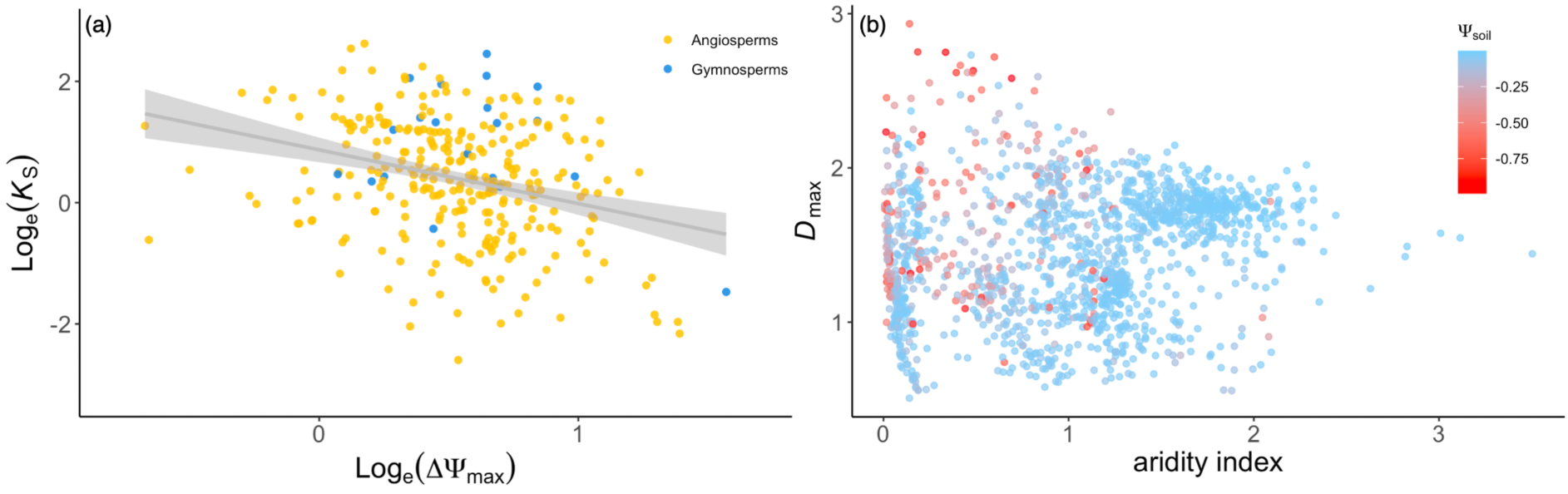
(a) The bivariate relationship between Log_e_-transformed hydraulic conductivity (*K*_S_) and maximum water potential difference between soil and leaf (ΔΨ_max_). The yellow dots are angiosperms and blue dots are gymnosperms. The gray line is fitted across all species. (b) The soil water potential (Ψ_soil_) variation along maximum vapour pressure deficit (*D*_max_) and aridity index for each species.

**Fig. S8.**
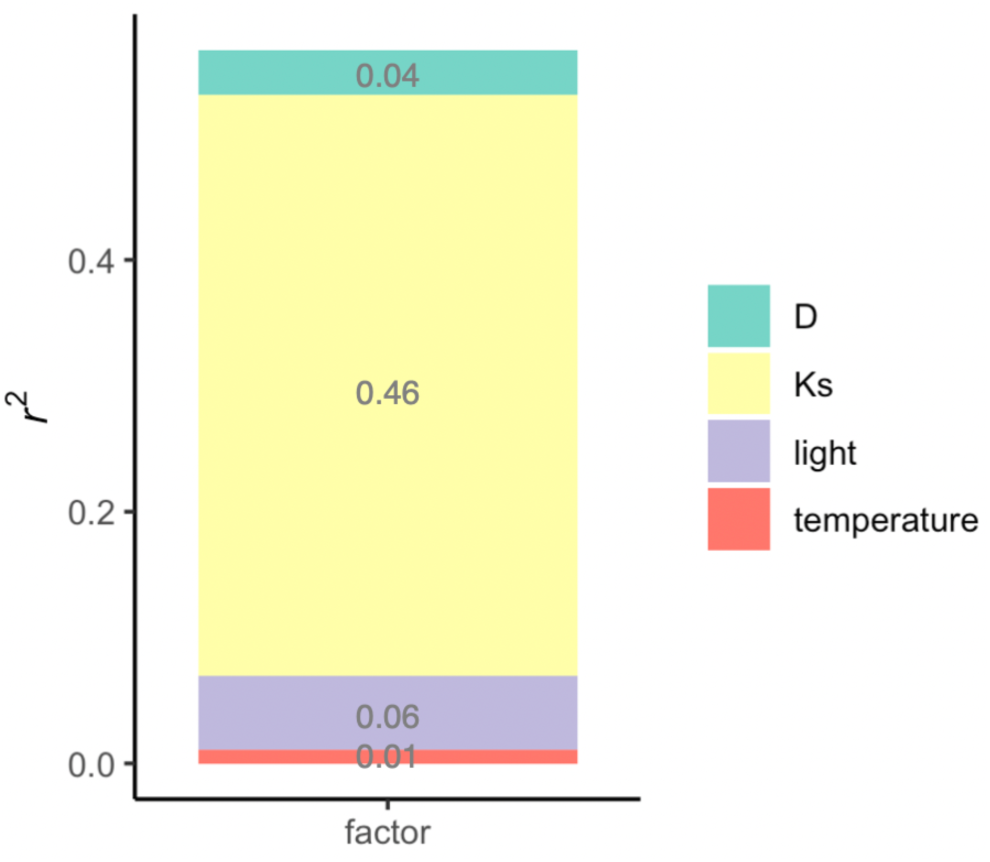
The contribution (*r*^2^) of different predictors to the ratio of sapwood to leaf area (*v*_H_) prediction. The green bar is vapour pressure deficit (*D*), yellow is hydraulic conductivity (*K*_S_), purple is irradiance/light and red is temperature.

**Fig. S9.**
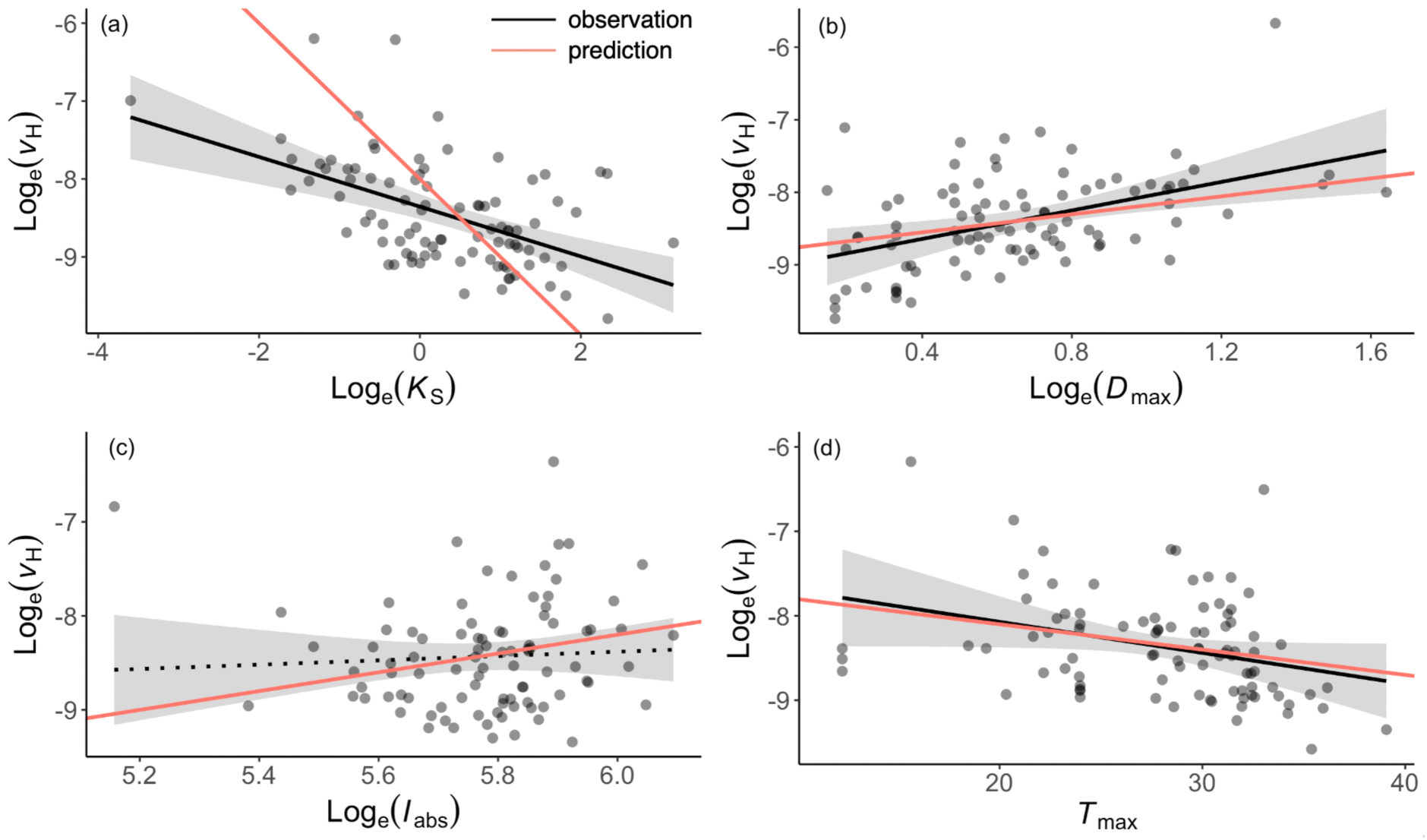
Partial residual plots from the multiple linear regression of log_e_-transformed the ratio of sapwood to leaf area (*v*_H_) against different predictors at site level using Dataset2. The predictors are shown in (a) sapwood-specific hydraulic conductivity (*K*_S_), (b) maximum vapour pressure deficit (*D*_max_), (c) mean irradiance (*I*_abs_), (d) maximum temperature (*T*_max_) during growing season. Black lines are the fitted across all sites and the gray shadings are the 95% confidence intervals around the fitted lines. The black solid lines are significant (*p*<0.05) and dotted line is insignificant (*p*>0.05). The red lines are theoretical sensitivities in equation (1).

**Fig. S10.**
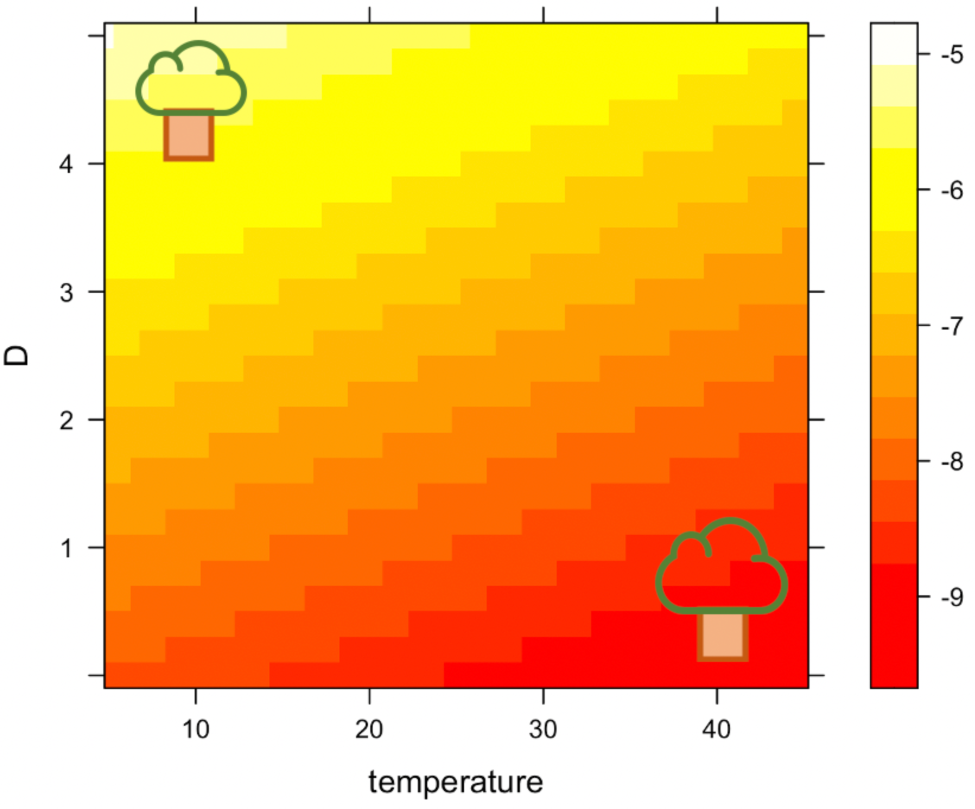
Theoretical variation of the ratio of sapwood to leaf area (*v*_H_) along temperature and vapour pressure deficit (*D*) with other predictors kept as median values.

## Notes

### Competing Interest Statement

The authors have declared no competing interest.

